# Interpretable neural networks prioritize cancer driver genes from genome-wide dependency landscapes

**DOI:** 10.1101/2025.04.28.651122

**Authors:** Qingyang Yin, Liang Chen

**Affiliations:** Department of Quantitative and Computational Biology, University of Southern California, Los Angeles, California, United States of America

## Abstract

Identifying cancer driver genes and their therapeutic impact remains a core challenge in computational cancer biology. We introduce xNNDriver and xAEDriver, two interpretable neural network frameworks that connect cancer mutations with genome-wide DepMap gene dependencies, pathway activity, and drug-response patterns. xNNDriver is a supervised pathway-guided model that evaluates whether a gene’s mutation status is encoded in the genome-wide dependency landscape; we interpret model fitness as a driver potential score, which quantifies the strength of this mutation-dependency signal and prioritizes genes with broad functional footprints. Across 3,008 candidate genes, xNNDriver recovers major established drivers and highlights literature-supported candidates, while pathway analyses reveal biologically coherent programs related to metabolism, growth factor signaling, and immune regulation. To capture combinatorial functional states, xAEDriver uses an unsupervised autoencoder to learn Driver Variant Representations (DVRs), latent binary features guided by the frequency distribution of known driver mutations. DVRs capture cell-line-specific dependency patterns and expression patterns and are associated with drug sensitivity and pathway activity. Together, these interpretable deep learning models demonstrate that gene dependency landscapes encode rich, interpretable signals of oncogenic function and provide a hypothesis-generating framework for prioritizing drivers, pathways, and therapeutic vulnerabilities for further experimental validation.

**Author summary:** Cancer is often driven by genetic changes that give tumor cells a growth advantage, but finding which changes matter and how they affect cell behavior remains difficult. In this study, we used large-scale gene-editing data from cancer cell lines to ask whether the pattern of genes a cell depends on can reveal information about its cancer-driving alterations. We developed interpretable neural-network models that connect mutation patterns, cell survival dependencies, and biological pathways. One model prioritizes genes whose mutation status leaves a clear functional signature across the cell. A second model summarizes broader, combined driver-like states in each cell line. These summaries were linked to known cancer biology, tissue type, and differences in drug response, suggesting that functional dependency data contain useful clues about cancer mechanisms. By connecting mutation patterns with functional dependencies and pathway activity, our approach helps identify candidate cancer drivers and interpretable drug-response patterns that can guide future experimental studies. Making the models interpretable allows researchers to move from large screening datasets toward biological explanations of cancer-driving processes.

## Introduction

One of the most critical and challenging areas in cancer biology is the identification of driver genes and mutations. Cancer driver mutations fuel tumor progression, unlike neutral passenger mutations (1). In recent years, machine learning and deep learning have shown great potential in bioinformatics, including applications for detecting cancer driver genes and mutations (2–6). However, current deep learning models predominantly utilize sequence data and three-dimensional protein structures to infer driver mutations (1), while functional genomics, especially gene editing, remains less explored. Additionally, the interpretability of these black-box models is often limited.

Conventional gene perturbation strategies, such as knockout and RNAi-mediated knockdown, enable researchers to study a gene’s loss-of-function effects on tumor progression, aiding in identifying cancer-driver genes and mutations (7). CRISPR-Cas9 has advanced gene perturbation in terms of precision and scale (8). The Cancer Dependency Map (DepMap) at the Broad Institute conducted genome-wide CRISPR-Cas9 knockout screens across a wide range of cancer cell lines (9, 10), providing gene dependency scores that reflect each gene’s importance for cell proliferation and survival in a cell-line-specific manner. While DepMap dependency scores provide valuable insights into the essentiality of genes in cancer cells, they are not sufficient on their own to conclusively determine whether a gene is a driver of cancer. Accurate identification of cancer driver genes requires an integrated analysis of dependency scores with other data types, such as DNA mutations, pathway involvement, and gene expression.

We hypothesize that the mutations with driver activity can leave detectable signatures in the genome-wide dependency landscape, whereas passenger mutations are less likely to produce coherent dependency shifts. To test the hypothesis, we built a supervised learning model to predict the mutation status (mutated or not) of each candidate gene, using the dependency scores of all screened genes in a cell line. The model’s fitness, expressed as the driver potential score of this candidate gene, prioritizes genes whose mutation status has a strong functional footprint in dependency data. We name this model **xNNDriver** (e**x**plainable **N**eural **N**etwork for **Driver** gene identification). Because the labels are derived from existing mutation annotations, xNNDriver is best viewed as a functional prioritization and interpretation framework rather than an independent de novo mutation-discovery tool. Our approach successfully identified both well-known driver genes and novel candidates. To discern the pathways through which these drivers contribute to tumorigenesis, we incorporated the Reactome pathway hierarchy (11) into the network architecture, designing each neuron to represent a distinct biological pathway.

To simultaneously infer multi-driver functional states in a cell line, we designed an interpretable autoencoder model that integrates both dependency scores and gene expression data. This unsupervised model is named **xAEDriver (**e**x**plainable **A**uto**E**ncoder for hidden overall **Driver** status identification). The latent layer of xAEDriver, which captures the key features of the input data in a compressed form, is guided by the mutation distribution of known driver SNPs. The latent representations are constrained to resemble the population-level distribution of driver mutations, but they are not forced to correspond to specific genes or variants. We refer to these latent representations as **D**river **V**ariant **R**epresentations (DVRs). xAEDriver is also interpretable, so it can reveal biological pathways that offer a deeper understanding of the driver mutation status. Notably, cell lines stratified by DVRs exhibited distinct drug responses, supporting the use of dependency-derived driver states to interpret treatment-relevant vulnerabilities.

## Results

### The fitness of the supervised xNNDriver quantifies cancer driver potential from dependency landscapes

Our supervised xNNDriver model links genome-wide gene dependency scores to the mutation status of a candidate driver gene in each cancer cell line. The mutation status of a true driver gene, but not passenger mutations, influences the impact of other genes’ knockout on tumor survival (dependency). In other words, the distinct genome-wide dependency scores can predict the mutation status of driver genes (high model fitness), but not those of passenger genes (low model fitness). We define the resulting model fitness as the driver potential score, a measure of mutation-dependency association and functional prioritization rather than direct proof of causality. A schematic overview of the workflow is shown in Fig 1A.

**Fig 1.**
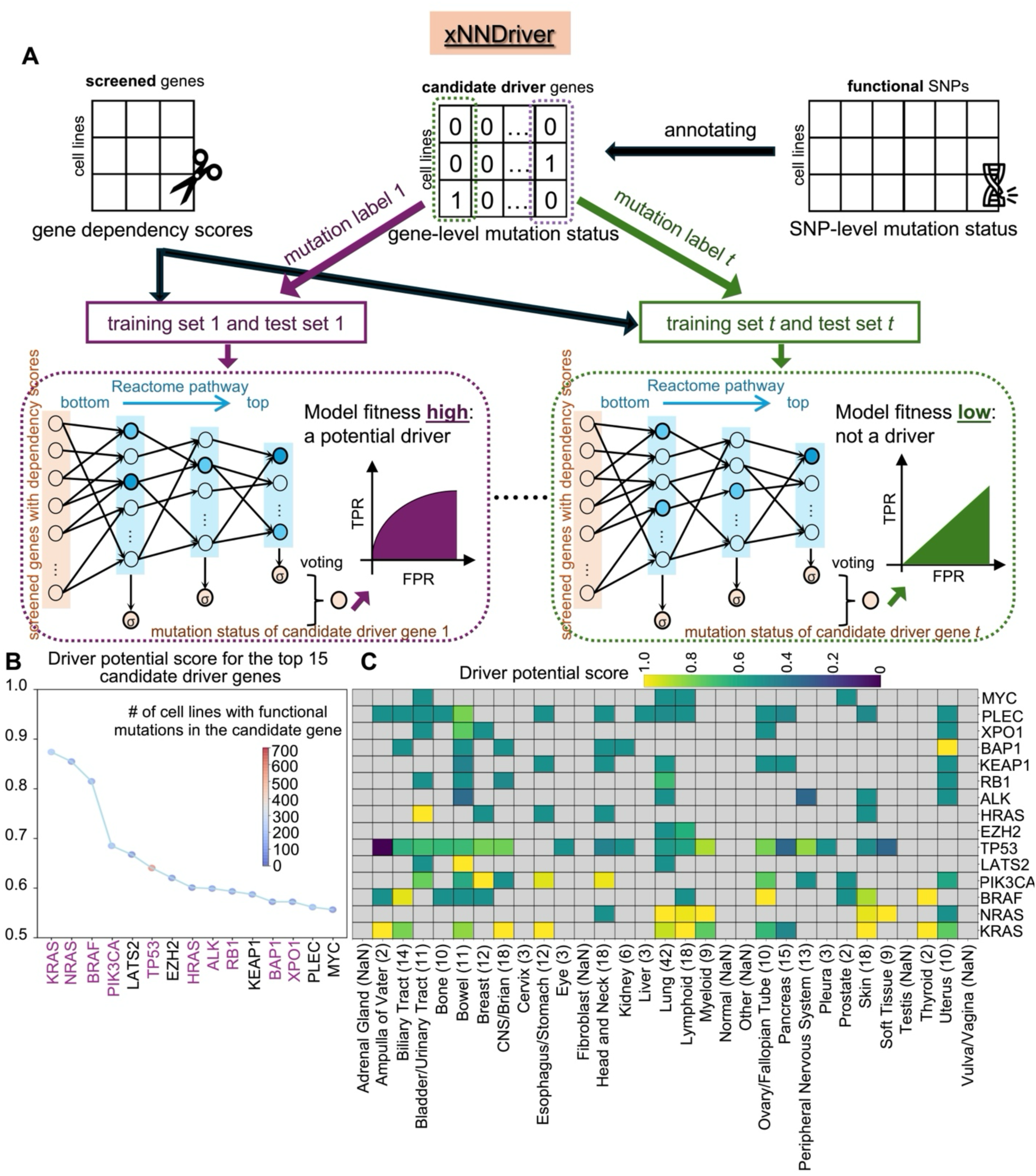
Supervised xNNDriver model for prioritizing individual candidate driver genes. **(A)** Workflow of xNNDriver. Functional SNPs are aggregated to annotate the gene-level mutation status for each candidate driver gene. Individual neural network models are trained using gene dependency scores as input features and candidate gene mutation status as labels. Data are split into training and test sets, and the model follows the Reactome pathway hierarchy. Predictions from all layers are integrated via voting. The model fitness is evaluated using the driver potential score, where a higher score indicates a stronger mutation-dependency association and higher priority as a candidate driver. **(B)** Prediction performance for the top 15 genes with the highest driver potential scores. Point color represents the number of cell lines with functional mutations in each candidate gene. A total of 1,095 cell lines (mutated and non-mutated) are considered for each model. Gene names are color-coded by driver annotation from DepMap (purple: driver; black: non-driver). **(C)** Tissue-level driver potential scores. The color of each square indicates the driver potential score within test cell lines from a given tissue, and the number following the tissue name denotes the count of test cell lines in that tissue.

To construct the training data, we determined each candidate gene’s mutation status based on the presence of functional SNP somatic mutations, as annotated by both the COSMIC (12) and ClinVar (13) databases. The model was trained using a biologically informed feedforward neural network whose architecture mirrors the pathway hierarchy acquired from the Reactome database. The input layer consists of screened genes with dependency scores, and the hidden layers represent pathways. The edges between the input layer and the first hidden layer capture pathway membership for each gene, while the edges between pathway layers represent the hierarchical relationships between pathway terms. Each pathway layer is followed by a predictive layer designed to predict the mutation status of a candidate driver gene. Predictions from all hidden layers are aggregated using a voting mechanism.

The model fitness of xNNDriver reflects the strength of association between a candidate gene’s mutation status and genome-wide dependency patterns. A high driver potential score therefore suggests that the gene’s mutation status is linked to a broad functional dependency signature in cancer cells and prioritizes it for driver-focused interpretation. Important pathways contributing to predictions are identified by contrasting the activation values of neural networks between mutated and non-mutated cell lines. We iteratively apply the xNNDriver model to all candidate genes (3,008 genes) to prioritize cancer driver genes and their associated important pathways.

### xNNDriver prioritizes established drivers and literature-supported candidate driver genes

We applied our supervised model, xNNDriver, to the 15% most frequently mutated genes in human cancers as determined by the CCLE project (14), totaling 3,008 candidate genes. Fig 1B illustrates the driver potential scores of the 15 highest-scoring genes, with complete results provided in S1 Table. Among the top 15 genes, ten are well-established cancer driver genes as documented in DepMap, such as *KRAS*, *NRAS*, *BRAF*, *PIK3CA, TP53*, and so on. This strong concordance supports that the dependency landscape contains functional information that captures core cancer driver biology.

The remaining genes, although not yet recognized as drivers in DepMap, have been suggested in the literature to possess driver-like functions. For example, Murakami et al. and Strazisar et al. (15, 16) reported that *LATS2* acts as a tumor suppressor essential for regulating the proliferation and/or survival of malignant mesothelioma cells. *EZH2* exhibits a dual role, functioning as either a tumor suppressor or an oncogene depending on the cellular context (17). Additionally, Sasaki et al. (18) identified *KEAP1* as an inhibitor of NRF2-induced cytoprotection and proposed it as a potential tumor suppressor. Collectively, these results demonstrate that xNNDriver can reprioritize known drivers and nominate literature-supported candidates for further validation.

To test the robustness of our model, we redefined mutation status using all SNP somatic mutations rather than only functional ones. The top 30 genes under this alternative annotation are shown in S1 Fig, with complete results in S2 Table. The recovery of major known drivers (*KRAS*, *NRAS*, *BRAF*, *PIK3CA*, *TP53*) and consistent identification of *EZH2* underscore the robustness of our approach.

We next examined tissue-specific driver activity. For the top 15 genes (Fig 1B), we recalculated driver potential scores within individual tissue types. As shown in Fig 1C, higher tissue-level scores indicate a gene’s stronger driver potential in that tissue. For instance, *BRAF* is strongly associated with thyroid cancers, consistent with the *BRAF V600E* mutation being the most common alteration in papillary thyroid carcinoma (19). Similarly, *PIK3CA* mutations frequently drive breast cancers (20). This tissue-resolved analysis highlights the context-specific roles of cancer drivers.

### xNNDriver reveals driver-associated pathway programs from dependency signatures

Using the interpretable xNNDriver model, we discovered important pathways associated with high-scoring candidate driver genes. Fig 2A shows the top 20 pathways, ranked by their importance measurements (*D*_*p*_ values), for the 15 genes with the highest driver potential scores. The complete list of pathways passing the *D*_*p*_ cutoff (default: 0.1) for these genes is provided in S3 Table. Metabolic pathways ranked prominently, including “metabolism,” “metabolism of proteins,” and “metabolism of lipids.” This aligns with Vander Heiden et al. (21), who demonstrated that altered cellular metabolism can drive tumor progression and inform cancer prognosis and therapy.

**Fig 2.**
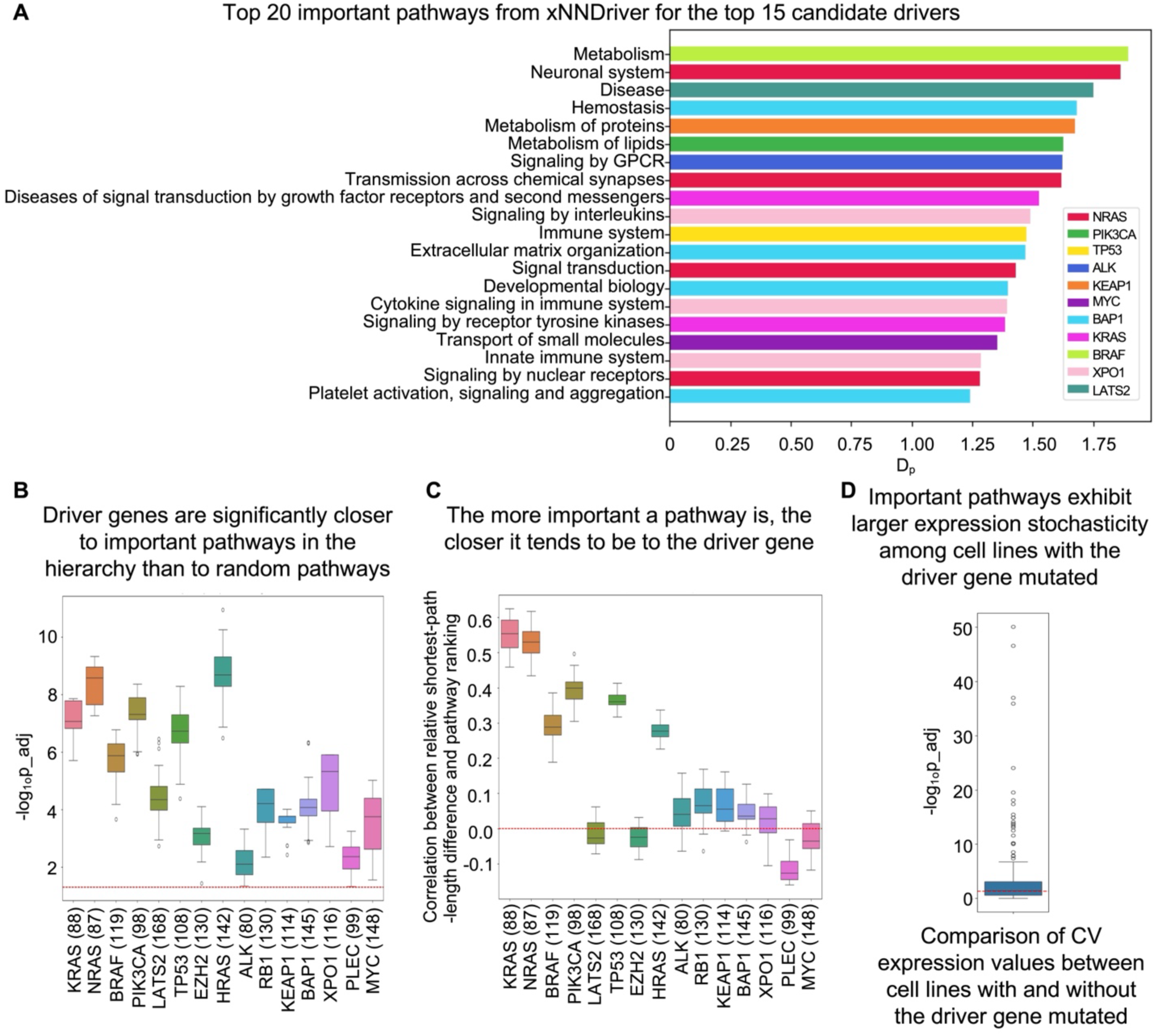
Analysis of important pathways from xNNDriver. **(A)** Top important pathways. The top 20 pathways are ranked by *D*_*p*_ values, aggregated from models of the 15 highest-scoring driver genes. For pathways identified in multiple models, the maximum *D*_*p*_ value is reported. **(B)** Pathway proximity to driver genes. Adjusted p-values are from the one-sided Wilcoxon signed-rank test comparing the shortest path lengths to the corresponding driver gene between identified and random pathways at the same hierarchy level. The random sampling procedures were repeated 20 times. Genes are sorted by their driver potential scores. The number in brackets next to the gene name indicates the number of important pathways identified from the corresponding gene model. The red dashed line marks the adjusted p-value cutoff of 0.05. **(C)** Correlation between pathway proximity and importance. Pearson correlation coefficients between relative shortest-path length differences and pathway rankings. The red dashed line marks the cutoff of 0. **(D)** Differential pathway expression variability. Adjusted p-values from one-sided Wilcoxon signed-rank tests comparing the expression CV values of all gene members of an important pathway between cell lines with and without the corresponding driver gene mutation are shown. The red dashed line marks the adjusted p-value cutoff of 0.05.

Additionally, signal transduction pathways, such as “diseases of signal transduction by growth factor receptors and second messengers,” were also highly ranked. This is consistent with Rho et al. (22), who highlighted growth factor signaling as a central mechanism in carcinogenesis. The “signaling by receptor tyrosine kinases” pathway governs essential processes in cell growth and survival, with mutations in key members (e.g., *EGFR*, *HER2*, *MET*) representing major cancer hallmarks (23, 24).

Interestingly, several immune-related pathways, such as “signaling by interleukins,” “cytokine signaling in the immune system,” and “innate immune system”, were identified as important pathways associated with the top 15 driver genes. These pathways are known to regulate inflammation, immune signaling, and tumor-immune interactions that contribute to cancer initiation and progression. Aberrant interleukin and cytokine signaling can activate oncogenic pathways such as JAK/STAT3 and PI3K-AKT, while dysregulation of innate immune signaling promotes chronic inflammation and immune evasion (25–27).

### Driver-associated pathways show Reactome proximity and independent expression variability

To assess the validity of the identified pathways, we measured their proximity to corresponding driver genes within the Reactome hierarchy. We calculated the shortest path length between each important pathway node (*D*_*p*_ ≥ 0.1) and its corresponding driver gene, comparing against random pathway nodes at the same level (20 random samplings). As shown in Fig 2B, important pathway nodes are significantly closer to their corresponding driver genes than random ones (Benjamini-Hochberg adjusted p-values < 0.05 for all comparisons, paired Wilcoxon signed-rank tests).

We further evaluated the relationship between pathway importance and hierarchical proximity. Across nearly all drivers, the relative difference in shortest-path length between important and random pathways positively correlated with pathway rank (Fig 2C). Thus, the more important a pathway is, the closer it is to the driver gene compared to random pathways at the same layer. Notably, genes with higher driver potential scores exhibit stronger correlations, as shown in Fig 2C, where genes are ordered by their scores. Together, these results support the use of *D*_*p*_ for interpretable pathway prioritization and reinforce the utility of the driver potential score as a functional association measure.

To explore the underlying mechanisms by which the pathways promote tumorigenesis, we examined pathway expression stochasticity. In our previous study (28), we found that pathways important for cell type stratification often exhibit differential expression stochasticity, measured by the coefficient of variation (CV; standard deviation over mean), rather than differences in mean expression between cell type groups. Here, for each important pathway, we calculated the expression CV for all genes within the pathway, comparing cell lines with the corresponding driver gene mutated versus unmutated, and assessed significance using a Wilcoxon signed-rank test. As shown in Fig 2D, of the 357 pathways analyzed (*D*_*p*_ ≥ 0.1 across the 15 highest-scoring gene models, S3 Table), 193 display a Benjamini-Hochberg adjusted p-value below 0.05. These findings emphasize that expression variability rather than mean differences may characterize driver-associated pathways, which is an insight often overlooked by traditional gene set enrichment analyses. Importantly, xNNDriver does not use expression data, so these stochasticity patterns are independent of model training and reflect intrinsic pathway characteristics.

### The unsupervised xAEDriver model learns multiple-driver functional representations

The success of xNNDriver suggests that genome-wide dependency scores for a tumor cell line contain information predictive of driver-associated functional states. Building on this, and recognizing that multiple driver mechanisms collectively shape tumorigenesis, we developed the unsupervised xAEDriver model to infer multiple driver variant representations (DVRs) that encapsulate cell-line-specific dependency and expression profiles while being regularized toward the population-level frequency distribution of known functional driver SNPs (Fig 3A). xAEDriver integrates gene dependency and expression data using an autoencoder with a fully connected encoder and a biologically informed decoder constrained by the Reactome pathway hierarchy. The latent layer captures lower-dimensional embeddings of the data. Here, to guide xAEDriver in generating hidden “driver mutation” information, we incorporated data from known functional driver SNPs to steer the autoencoder’s training, ensuring that the lower-dimensional embeddings (referred to as DVRs) have frequency distributions similar to real driver SNPs (Materials and methods). We then applied layer-wise relevance propagation (LRP) to identify important pathways contributing to the reconstruction, enabling us to uncover additional biological insights related to the latent functional states.

**Fig 3.**
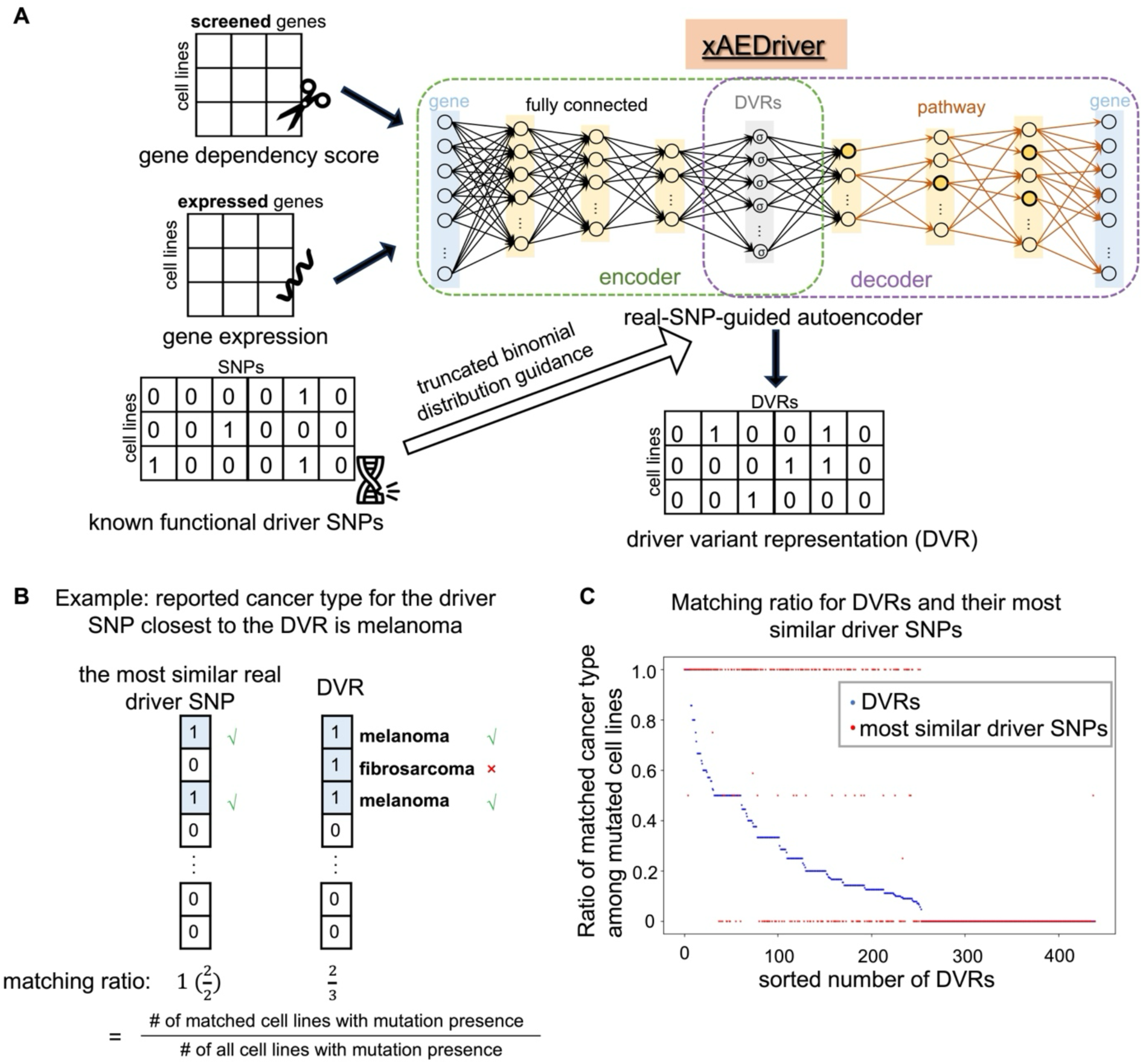
Unsupervised xAEDriver model to reconstruct driver variant representations (DVRs). **(A)** Workflow of xAEDriver. Gene dependency and expression data are jointly used to train an autoencoder with a fully connected encoder and a biologically constrained decoder aligned with the Reactome hierarchy. The encoder produces low-dimensional embeddings (DVRs) representing cell-line-specific driver states. Known driver SNPs are incorporated into training using a truncated binomial modeling approach**. (B)** Illustration of cancer type matching ratio. As an example, among three cell lines with mutations of the DVR (value = 1), two match the reported cancer type (melanoma) for the corresponding (the most similar) known driver SNP, yielding a matching ratio of 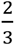. Similarly, for the most similar real driver SNP, the matching ratio is 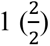. **(C)** Comparison of cancer-type matching ratios between DVRs (blue) and their most similar known driver SNPs (red). The data are sorted in descending order based on the matching ratios of the DVRs.

### DVRs recapitulate driver-mutation frequency and represent the overall driver mutation status of a cancer cell line

To evaluate the effectiveness of the guided learning in xAEDriver, we compared the mutation distribution of known driver SNPs with that of DVRs across all cell lines (S2 Fig). The mutation distribution of DVRs closely matched that of actual driver mutations, as demonstrated by the Earth Mover’s Distance (EMD) of 5.4 (on a 0-1,000 scale) between their mutation patterns. EMD, a metric for assessing distribution similarity, represents the minimum effort required to transform one distribution into another (29). These results underscore the effectiveness of the regularization term in mimicking the distribution of real mutations, guiding xAEDriver to actually learn DVRs in the hidden layer.

Next, we examined whether each DVR’s artificial mutation status (coded as 1 in the artificial representation vector) matched the reported cancer type of its closest real SNP, determined via Euclidean distance. The cancer type matching ratio was calculated as illustrated in Fig 3B. After excluding 481 DVRs with non-unique matched real SNPs, 61 out of 440 DVRs achieved a matching ratio ≥ 0.5 (Fig 3C), suggesting that many DVRs capture cancer-type-specific mutation patterns. Among the remaining 379 DVRs with a matching ratio below 0.5, 263 (or 69%) still had a matching ratio greater than or equal to that of their corresponding real SNP. This suggests that the associated real mutations are often observed in a cancer type different from the one reported. This could be due to several factors: the driver mutation may drive tumorigenesis in another cellular context or a different cancer type, or the mutation may emerge during cancer progression as a compensatory mechanism or an accelerating “second hit”.

### xAEDriver highlights pathways central to tumor progression

Using layer-wise relevance propagation (LRP) in xAEDriver, we computed pathway relevance scores (*R*_*p*,*c*_) for each pathway *p* and cell line *c*, where higher values indicate greater contribution to the reconstruction of the autoencoder. Averaging across all cell lines yielded overall pathway importance (*R*_*p*_). Fig 4 shows the top 20 pathways ranked by *R*_*p*_ values, with the complete list provided in S4 Table. The top pathways, such as “complement cascade” (rank 1), “TNFR2 non-canonical NF-kB pathway” (rank 2), and “HuR (ELAVL1) binds and stabilizes mRNA” (rank 6), have been previously linked to tumor progression. For example, Rutkowski et al. (30) highlighted the role of the complement cascade in driving cancer by promoting key hallmarks of its progression. Albini et al. (31) demonstrated that *RAF2* inactivating mutations increase alternative NF-kB pathway activation in mantle cell lymphoma, diffuse large B-cell lymphoma, and multiple myeloma. Cai et al. (32) revealed that *ELAVL1* is an m6A-binding protein highly expressed in multiple tumors that regulates RNA stability to promote tumor progression. These results confirm that xAEDriver effectively uncovers biologically meaningful pathways central to tumor biology.

**Fig 4.**
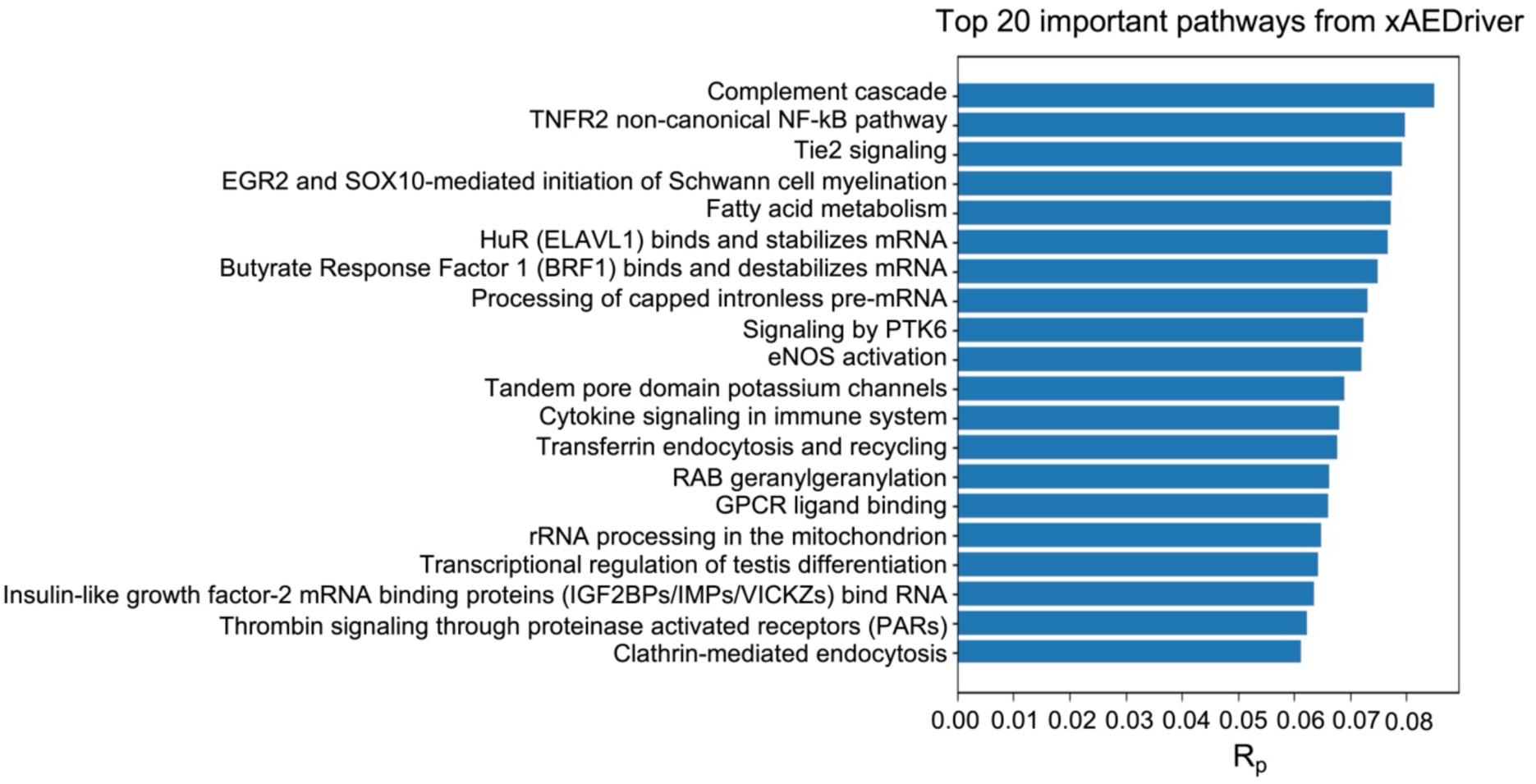
Important pathways identified from xAEDriver. The top 20 pathways ordered by *R*_*p*_ values are shown.

### DVRs link multi-driver functional states to drug response

Having established that xAEDriver captures meaningful driver variant representations (DVRs), we next investigated whether these DVRs are associated with functional outcomes in cancer cells, specifically their ability to reflect tissue-specific drug sensitivity patterns and provide mechanistic insights into differential drug responses. To address this, we constructed LASSO regression models to predict drug sensitivity (AUC) for each drug using the DVR binary matrix across our panel of cancer cell lines (Materials and methods). DVR-based models significantly outperformed those trained on an equivalent number of random somatic mutations in the test set. Twelve DVR models achieved a test-set *R*^2^ > 0.05, whereas none of the 50 random models met this threshold (complete list in S5 Table).

For each of the twelve DVR-based models passing this performance threshold, we identified the ten most influential DVRs based on the absolute values of their LASSO coefficients, comparing them to a control set of ten randomly selected DVRs. The biological relevance of these top DVRs was evaluated using a tissue concordance score (Materials and methods), which quantifies the alignment between each DVR’s mutation distribution across tissues and the tissue distribution of the most drug-sensitive cell lines (lowest AUC values). These top DVRs showed significantly higher tissue concordance scores than random controls, indicating that they capture tissue-specific drug response patterns (p = 0.0005, paired Wilcoxon signed-rank test, Fig 5A). Repeating the random sampling ten times produced a consistent average p-value of 0.0015. These results demonstrate that DVRs effectively prioritize biologically meaningful, tissue-specific features associated with drug sensitivity.

**Fig 5.**
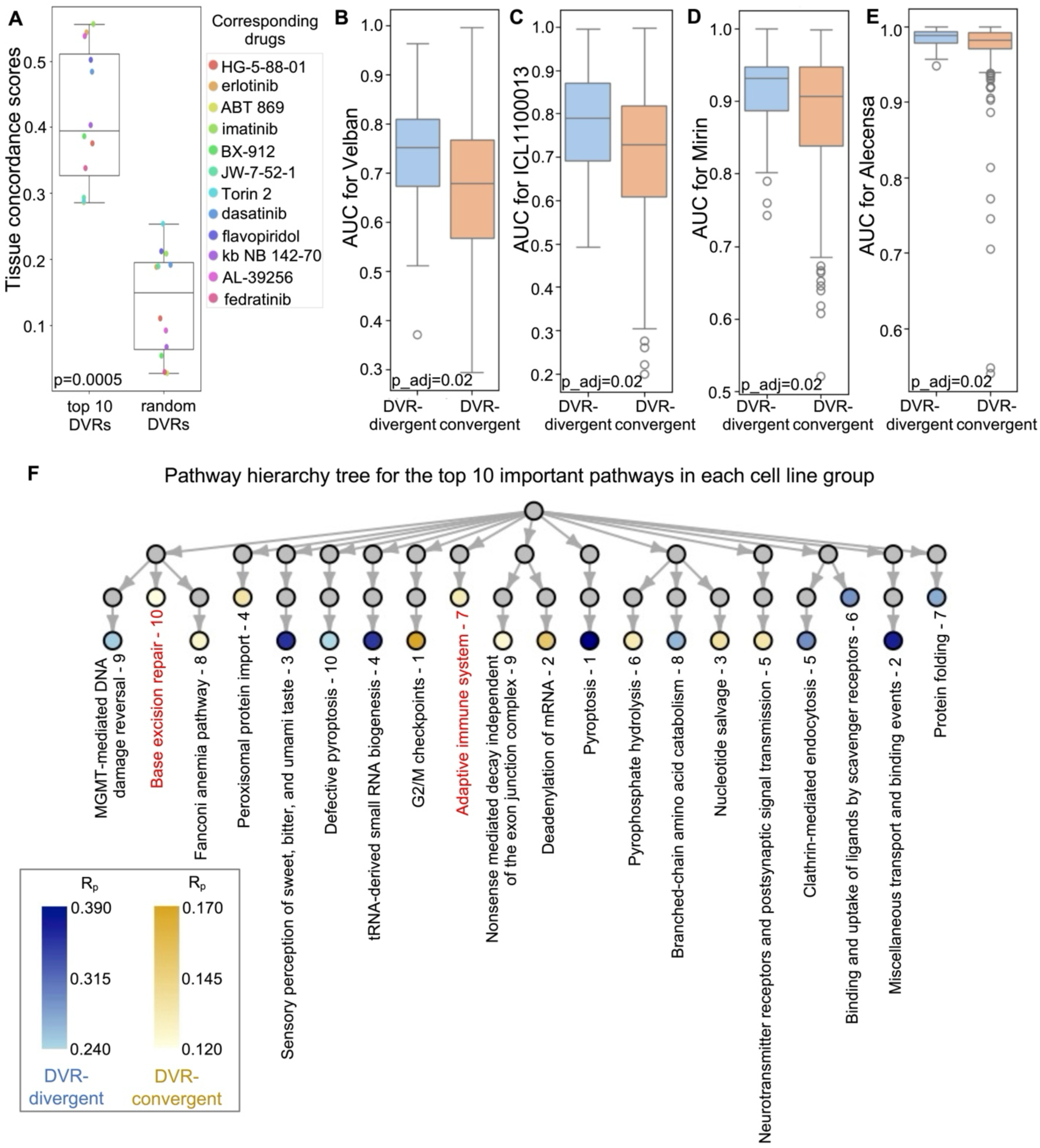
DVRs from xAEDriver reveal drug-response patterns via LASSO regression and distance-based stratification. **(A)** Differential tissue concordance of DVRs. The top 10 DVRs from LASSO models for twelve LASSO-explainable drugs show significantly higher tissue concordance scores than random DVRs (p = 0.0005, paired Wilcoxon signed-rank test). **(B-E)** Differential drug sensitivity. Cell lines stratified by DVR dissimilarity, DVR-divergent (high dissimilarity) and DVR-convergent (low dissimilarity), exhibit distinct drug responses. The DVR-convergent group shows significantly higher sensitivity (lower AUC) to drugs **(B)** Velban, **(C)** ICL1100013, **(D)** Mirin, and **(E)** Alecensa (all adjusted p = 0.02, t-tests). **(F)** Pathway hierarchy tree for the top 10 important pathways in the DVR-divergent and DVR-convergent groups. Highlighted pathways illustrate representative differences in *R*_*p*_(pathway importance) between groups. Gray nodes denote parent pathways connecting these top-ranked nodes, while non-relevant child and sibling nodes are omitted for clarity.

Next, we stratified cancer cell lines based on their DVR binary matrix using a distance-based approach (Materials and methods). This analysis divided the cell lines into a DVR-divergent (high dissimilarity) group (n=76) and a DVR-convergent (low dissimilarity) group (n=430). Comparison of drug responses between these two groups identified seven drugs with significantly different AUC values (Benjamini-Hochberg adjusted p-value < 0.05, t-test). Fig 5B-E highlight four representative drugs, including Velban, ICL1100013, Mirin, and Alecensa. Interestingly, all four drugs were more effective in the larger DVR-convergent group, suggesting shared vulnerabilities across a broad spectrum of cancer cell lines. The remaining three drugs with significant differences were SN38, MPS-1-IN-1, and Y-39983.

To further assess the relationship between DVR and drug response, we correlated the pairwise Euclidean distances between cell lines in the DVR space with the pairwise Pearson’s correlations of their drug response (AUC) profiles. This analysis revealed a significant, though modest, negative correlation (PCC = −0.04, p-value = 5.5×10^-39^), indicating that cell lines with greater DVR dissimilarity tend to exhibit less similar drug sensitivity patterns. These results demonstrate that DVR-based stratification can effectively distinguish cell lines with distinct drug response profiles.

To explore the biological mechanisms underlying these differences, we analyzed pathway relevance within each cell line group. All 680 pathways were ranked by their average relevance scores across cell lines (S6 Table), revealing group-specific enrichment among the top pathways (Fig 5F). For example, the “Adaptive immune system” pathway ranked 7th in the DVR-convergent group but 654th in the DVR-divergent group, consistent with higher sensitivity to Velban and ICL1100013 in the DVR-convergent group (Fig 5B-C). Previous findings show that local low-dose Velban injection enhances adaptive immunity via dendritic cell maturation (33), while ICL1100013 targets *NMT1*, an enzyme critical for T-cell receptor signaling and immune function (34, 35). Similarly, the “Base excision repair” (BER) pathway ranked much higher in the DVR-convergent group (10th vs. 564th in the DVR-divergent group), consistent with increased sensitivity to Mirin (an *MRE11* inhibitor) and Alecensa (an *ALK* inhibitor) (Fig 5D-E). These associations are mechanistically coherent: *MRE11* plays key roles in BER and single-strand break repair (36), while Alecensa resistance, characterized by low *SFTPD* expression, has been linked to elevated BER pathway activity (37). Together, these findings suggest that enhanced immune and DNA repair pathway activities underlie the heightened drug sensitivity observed in the DVR-convergent group.

Overall, these results demonstrate that DVR-based representations capture biologically and therapeutically relevant signals, enabling the identification of tissue-specific patterns, drug sensitivities, and underlying pathway mechanisms in cancer cell lines.

## Discussion

In this study, we introduce interpretable functional-genomics frameworks for linking cancer driver alterations to genome-wide dependency landscapes, pathway programs, and drug-response structure in cancer cell lines. xNNDriver prioritizes candidate driver genes by quantifying how strongly their mutation status is reflected in dependency profiles, while xAEDriver learns multi-driver functional representations that summarize broader oncogenic states. Together, these models show that DepMap dependency data encode biologically interpretable signals of driver activity and treatment-relevant vulnerability.

xNNDriver reframes driver identification from a purely mutation-frequency paradigm to a systems-level functional consequence paradigm. The fact that known drivers score highly supports the biological signal embedded in dependency landscapes. Furthermore, the model is applied uniformly to 3,008 candidate genes and identifies both established and literature-supported but not yet DepMap-annotated candidates. Importantly, the driver potential score quantifies the functional footprint of mutations rather than their recurrence alone.

Given that the dependency score of the candidate driver gene provides a quantifiable functional signature of driver activity, it was included as an input feature and could be intrinsically selected during model training. For example, loss of an oncogene often reduces cell viability (low dependency score), whereas loss of a tumor suppressor enhances growth (high dependency score). As shown in S3 Figure, *TP53*, a tumor suppressor, exhibits the highest mean dependency score (0.426), while *NRAS*, an oncogene, shows a low value (−0.654). However, *HRAS*, another oncogenic driver, displays a near-neutral score (−0.107), underscoring that single-gene dependency alone provides only partial information about driver status.

To model complex combinatorial functional states, we developed the unsupervised xAEDriver model. While xNNDriver excels at evaluating individual driver genes, it is not inherently designed to capture systemic, multi-driver states. In contrast, xAEDriver represents a conceptual shift from single-driver evaluation to multi-driver functional state inference by encoding cell-line-specific patterns that mimic the behavior of real functional driver variants through guided learning.

DVRs showed biologically interpretable and drug-associated patterns. Using LASSO regression, we demonstrated that DVRs’ predictive of drug response also exhibited strong tissue-type specificity, supporting their biological relevance. Furthermore, DVR-based stratification of cancer cell lines revealed distinct groups with significantly different drug sensitivity profiles and pathway activities. This stratification identified specific vulnerabilities, such as enhanced response to immune-related or DNA repair-targeting drugs, highlighting the translational potential of DVRs in predicting therapeutic responses and informing precision oncology strategies.

To enhance interpretability, both xNNDriver and xAEDriver incorporated biologically informed neural network architectures guided by the Reactome pathway hierarchy. Rather than acting merely as a sparsity constraint, this design embeds the hierarchy directly into the network to aggregate signals across levels, enabling the mechanistic attribution of driver effects. We do not view the pathway-constrained architecture alone as proof of a causal mechanism; instead, pathway attributions are supported by standalone proximity analysis, expression-stochasticity analyses, and consistency with external literature. In xNNDriver, the identified important pathways demonstrate hierarchical coherence and expression variability patterns. In xAEDriver, a biologically constrained decoder preserves pathway structure while allowing the encoder to learn flexible representations, since fully constraining both sides proved too restrictive and often produced trivial DVRs. Ultimately, the pathways identified by both models confirm the biological robustness of our framework, providing interpretable and biologically grounded insights beyond conventional pathway enrichment analyses.

Looking ahead, expanding the models to include larger and more diverse datasets will improve the detection of both common and rare driver mutations across different cancer types. This broader scope will refine our understanding of gene dependencies and mutation synergies, leading to more accurate prediction of therapeutic vulnerabilities. Ultimately, the integration of dependency data with mutational and pharmacological information will advance precision oncology by enabling more personalized and effective cancer treatment strategies.

## Materials and methods

### Datasets

#### The DepMap database

We utilized data from the Cancer Dependency Map (DepMap) projects (23Q2 version). This release contains data from CRISPR-Cas9 knockout screens across multiple tumor types, as part of Broad’s Achilles (38) and Sanger’s SCORE projects (39). It also includes genomic and transcriptomic characterizations from the CCLE project (40).

The primary dataset includes CRISPR-Cas9 essentiality screens for 17,931 genes across 1,095 cell lines (9, 10), resulting in a cell-line-by-screened-gene matrix. Each entry represents the “dependency score” of a screened gene, which was processed from the log-fold change in sgRNA read counts before and after knockout. A lower dependency score indicates a stronger reduction in cell viability following gene knockout, implying that the gene is essential for cell survival. Quality control and normalization were performed by DepMap.

Somatic point mutations and indels for the DepMap cell lines were obtained from the same source (41), totalling 1,408,099 mutations, including 1,290,926 single-nucleotide polymorphisms (SNPs). Gene expression data (RNA-seq) were also retrieved as transcripts per million (TPM) for 19,194 genes across 1,450 cell lines, log-transformed with a pseudo-count of one. A total of 1,019 cell lines had both dependency score and gene expression data available.

We also downloaded drug sensitivity data for the cell lines (42). From the raw data, fold-change viability values were calculated using the R package gdscIC50 (43), and standard four-parameter monotonic log-logistic dose-response curves were fitted using the R package dr4pl (44). Drug sensitivity was then quantified as the area under the curve (AUC), covering 320 drugs across 962 cell lines. Among these, 506 cell lines also had dependency scores and gene expression data available.

#### The COSMIC database

We obtained mutation data from the Catalogue of Somatic Mutations in Cancer (COSMIC) Cancer Mutation Census (12), filtering for single-nucleotide variants (SNVs). To identify functionally significant mutations, we retained those annotated as “high” or “moderate” impact based on consequence predictions from the Ensembl Variation database (45).

#### The ClinVar database

ClinVar compiles relationships between human genetic variants and phenotypic outcomes (13). Although primarily focused on germline mutations, it includes somatic variants as well. We downloaded the ClinVar summary table and filtered for SNVs annotated as: pathogenic, likely pathogenic, affects, drug response, or confers sensitivity.

We merged the filtered COSMIC and ClinVar SNVs, yielding 28,459 clinically significant variants also documented in DepMap. Among these, 340 were located in known driver genes with mutations observed in at least one cell line, which we designated as known driver mutations for model training and validation.

### xNNDriver: Supervised model for identifying driver genes

#### Training and test data

Gene names from the considered datasets were converted to Ensemble IDs using the DAVID portal (46), ensuring consistency with the Reactome database. Genes absent from the Reactome database were excluded. Dependency scores were split into training (75%) and test (25%) sets, with five-fold cross-validation on the training set for hyperparameter tuning. Because mutation-positive samples were much fewer than negatives, we balanced the classes during the training process by random upsampling of positive samples to achieve a 1:1 ratio.

#### Pathway hierarchy

The Reactome database (11) was used to construct a hierarchical pathway tree using the NetworkX Python library. Internal nodes represent pathways, and leaves correspond to genes. To simplify the hierarchy and reduce redundancy, each gene was linked only to its closest pathway nodes. A hyperparameter *l* controlled the number of pathway levels included. Genes or pathways absent from the data were excluded. The resulting structure was encoded as a binary mask matrix *M*, where *M*_*i,j*_ = 1 denotes a valid connection. Following our previous work (28), multi-layer predictive designs were employed to enhance model accuracy.

#### Neural network construction

We built a feedforward neural network based on the mask matrix. The input layer consists of CRISPR-Cas9 screened genes, with their corresponding dependency scores as input data. The size of the input layer corresponds to the number of genes allocated to the lowest level of the pathway hierarchy. For each layer, the input is denoted as *x* and the output as *y*. The output of each layer is calculated as: *y* = *f*[(*MW*)*x* + *b*], where *f* is the activation function (tanh), *M* is the mask matrix, *W* is the weight matrix, and *b* is the bias vector. Here, *MW* is the element-wise multiplication (Hadamard product) of *M* and *W*. Binary predictions were generated using a sigmoid function: 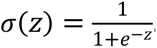. A cell line was classified as 1 (mutated) if *σ* > 0.5. Training used the Adam optimizer, cross-entropy loss, and L_2_ regularization. Final predictions were aggregated across all predictive layers via majority voting, and model performance was quantified by the AUC metric on the test set, referred to as the driver potential score.

#### Hyperparameter tuning

We applied stepwise hyperparameter tuning with five-fold cross-validation on the training set of the model for the candidate gene *KRAS* to optimize hyperparameters. Hyperparameters were tuned sequentially in the following order: the number of levels in the pathway hierarchy tree (*t*), L_2_ regularization hyperparameter, learning rate, batch size, and number of epochs. The optimal hyperparameter values based on the highest AUC were *l* = 3, regularization = 0.0001, learning rate = 0.01, batch size = 128, and epochs = 100.

#### Identification of important pathways in xNNDriver

Pathway importance was measured as the absolute difference in activation values between cell lines with and without mutation for a given pathway *p*: *d*_*p*_ = |*A*_*p*,0_ − *A*_*p*,1_|. If a pathway appears in the paths of multiple prediction layers (nested subnetworks), we chose the maximum value of *d*_*p*_ across all paths as the final importance measure, denoted as *D*_*p*_. Pathways with a *D*_*p*_ value greater than a predefined threshold (default: 0.1) were considered important. If the pathway node was an artificial node introduced to preserve the pathway tree structure, the original pathway node was retrieved.

#### Proximity analysis in the Reactome hierarchy

For each candidate driver gene, we calculated the shortest path length between its important pathways and the gene within the Reactome tree using NetworkX. Random pathways from the same hierarchical level served as controls. Statistical significance was assessed using paired Wilcoxon signed-rank tests. Correlations between relative path-length differences (the shortest path length for an important node minus that for a random pathway node, divided by the former) and *D*_*p*_ ranks (sorted descendingly) were computed using Pearson correlation coefficients across 20 randomizations.

#### Expression stochasticity of the pathways

Pathway expression variability was quantified by the coefficient of variation (CV: standard deviation over mean) of member genes across cell lines with or without the candidate gene mutation. Differences in CV distributions were tested using the Wilcoxon signed-rank test.

### xAEDriver: Unsupervised model for multi-driver representation learning

#### Training data preparation

The model xAEDriver utilized both gene dependency score and gene expression data as input features. Cell lines present in only one of the two data sources were excluded. For each gene *j* in cell line *i*, each feature was normalized within genes: 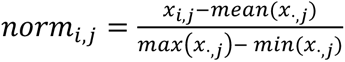 normalization, the gene dependency and expression data were merged into a unified matrix by aligning cell lines.

#### Guided learning using known driver SNPs

Known driver SNP mutation frequencies across cell lines were modeled using a truncated binomial distribution to exclude both endpoints. Let *X* represent the number of cell lines in which a known driver SNP is present as the mutated allele. The truncated binomial distribution for *X* is:

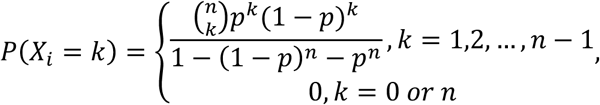

where *n* (=1,019) is the number of cell lines, and *p* is the unknown probability of SNP mutation being present in the population. Note that for all known driver SNPs, *k* is neither 0 nor *n*. Parameter *p* was estimated via maximum likelihood from the 340 known driver SNPs.

#### Autoencoder construction

The autoencoder consists of a fully connected encoder and a biologically informed decoder. The encoder and decoder share a symmetric structure in terms of node numbers at each layer, and both use the tanh activation function. However, in the decoder, certain neuron connections are masked according to the pathway-gene mask matrix, while the encoder remains fully connected.

The pathway-gene mask matrix was constructed following the approach outlined in the description for xNNDriver. The key distinction was in how we handled genes that appeared in both the dependency score and expression datasets. For such genes, the pathway was linked to both the “dependency” node and the “expression” node, and both entries in the mask matrix were set to one. For genes present in only one dataset, connections were handled as described before.

The lower-dimensional representation, referred to as driver variant representations (DVRs), is produced in the latent layer, which consists of 1024 neurons. Each DVR is represented by a neuron in the latent layer with values constrained to 0 or 1, indicating the mutation presence or absence of the DVR. The sigmoid activation function is used in the latent layer, ensuring the binary nature of the DVRs (rounded to 0 or 1). The autoencoder was trained by the Adam optimizer, and the cost function includes both reconstruction loss (measured as the mean squared error between the output layer and the input layer) and regularization terms, which include an L_2_ regularization term and a binomial distribution penalty.

Specifically, the binomial distribution penalty term encourages the DVRs to resemble real driver SNPs. In each epoch, after obtaining the lower-dimensional representations, we computed the sum for each DVR neuron across all cell lines and denoted it as *x*_*i*_. The binomial distribution penalty term is defined as the negative logarithmic likelihood across all DVRs (q=1,024), calculated as:

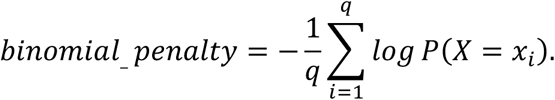

If *P*(*X* = *x*_*i*_) is equal to 0, the training process will be terminated, and the rounded DVRs from the previous epoch will be used as the final output.

#### Hyperparameter tuning

Grid search was used to tune the model’s hyperparameters. For each hyperparameter combination, we recorded the average loss over five repeats and selected the combination yielding the lowest loss as the default. The default values for the L_2_ regularization hyperparameter and binomial distribution penalty hyperparameters were set to 0.1 and 0.001, respectively. The default learning rate was 0.01. The default number of epochs was 1,000. The batch size was fixed at 1,019, corresponding to the total number of cell lines, as it is required to remain constant to compute the binomial distribution penalty.

#### Measuring pathway importance by layer-wise relevance propagation

To assess the contribution of pathways in generating DVRs, we employed layer-wise relevance propagation (LRP) with the epsilon rule (47). LRP decomposes the prediction by propagating relevance scores backward through the autoencoder’s layers. For a given cell line *c*, the relevance score *r*_j,*c*_ for neuron *j* is calculated as:

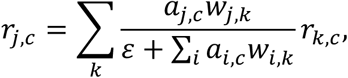

where *k* denotes a neuron in the subsequent layer (closer to the output), and *i* iterates over all neurons in the same layer as *j* that connect to *k*. Given the pathway-constrained hierarchy of our architecture, these connections are sparse, meaning *w*_*i*,*k*_ is non-zero only for biologically established links. Here, *a*_j,*c*_ and *a*_*i*,*c*_ represent the forward-pass activation values, *w*_j,*k*_ and *w*_*i*,*k*_ are the corresponding connection weights, and *ε* is a small positive constant. The relevance scores are propagated from the output layer to the input layer. We compute normalized relevance scores for each layer using z-scores, and the pathway importance is quantified by examining the decoder layer. The final normalized relevance score for pathway *j* and cell line *c* is denoted as *R*_j,*c*_.

#### LASSO regression model for DVR-drug association

For each drug, a LASSO regression model was built with DVRs as predictors and drug AUC values as responses. Data were split into training (80%) and test (20%) sets. Models were trained using a LASSO regularization parameter (α) of 0.001 and a maximum of 2,000 iterations. Predictive models (*R²* > 0.05) were analyzed for their top 10 DVRs based on coefficient magnitude. LASSO models were also built on 50 sets of random mutation controls for comparison.

To evaluate the biological relevance of the top DVRs, we calculated a tissue concordance score for each DVR. For a given drug, we defined two groups of cell lines: (1) the DVR-positive group, consisting of all cell lines in which the DVR was present (value = 1), and (2) the drug-sensitive group, consisting of an equal number of cell lines showing the strongest response to the drug (i.e., the lowest AUC values). The tissue concordance score was defined as the degree of overlap in tissue-type composition between these two groups, normalized by group size. This score reflects the degree of tissue-specific association between DVR presence and drug response.

As a control, for each drug, we randomly selected ten DVRs that were not among the top ten and repeated the concordance analysis. We then compared the tissue concordance scores between the top DVRs and the random controls using a paired Wilcoxon signed-rank test to assess statistical significance.

#### Distance-based DVR stratification for drug sensitivity patterns

We calculated the total Euclidean distance of each cell line to all others in DVR space and stratified lines into DVR-divergent (large total distance) and DVR-convergent (small total distance) groups using the 85th percentile cutoff. Drug sensitivity (AUC) differences were assessed with Welch’s t-test, omitting cell lines lacking drug data.

## Supporting information

**S1 Fig. Prediction performance of xNNDriver for the top 30 genes with the highest driver potential scores using all SNPs.** All somatic SNP mutations, no matter whether annotated as functional or not, are used. Data point color indicates the number of cell lines with mutations in that gene. Gene name color distinguishes whether a gene is a known driver or not (purple: driver, black: non-driver), as documented by DepMap.

**S2 Fig. Comparison of mutation frequency distributions between DVRs from xAEDriver and known functional driver SNPs**. The histograms display the frequency of mutation occurrences across cell lines for (**A**) DVRs and (**B**) known driver SNPs.

**S3 Fig. Distribution of mean dependency scores of all genes.** The histogram illustrates the distribution of mean dependency scores across all cell lines for all genes in the DepMap gene dependency dataset.

**S1 Table. Comprehensive results of all xNNDriver models using functional SNPs.** Each model corresponds to a frequently mutated candidate gene (3,008 in total). Information includes the following metrics: driver potential score, the number of cell lines with mutation presence, the number of cell lines without mutation presence, and whether the gene is annotated as a driver based on DepMap.

**S2 Table. Comprehensive results of all xNNDriver models using all SNPs.** Each model corresponds to a frequently mutated candidate gene (3,008 in total). Information includes the following metrics: driver potential score, the number of cell lines with mutation presence, the number of cell lines without mutation presence, and whether the gene is a driver based on DepMap.

**S3 Table. Important pathways identified from xNNDriver for candidate driver genes.** This table includes a comprehensive summary combining results for all 15 genes, as well as 15 individual sheets detailing results for each gene. The overall table also provides the adjusted p-values from the Wilcoxon tests on the comparison of coefficients of variation of pathway expression between cell lines with and without mutations.

**S4 Table. Pathway importance from xAEDriver.** This table includes the average *R*_*p*_ values across all cell lines. Pathways marked with “_copy1” indicate artificial nodes created to preserve the pathway tree structure.

**S5 Table. Test set** *R*^*2*^**values of LASSO regression models based on DVRs and random somatic mutations.** Random control results are averaged over 50 runs.

**S6 Table. Average pathway importance (***R*_*p*_ **values) and rankings for the DVR-divergent and DVR-convergent groups.** Reported metrics include average *R*_*p*_ and rank for each group.

## Acknowledgments

We thank Dr. Sika Zheng at the University of California, Riverside, Yingtong Liu, and all members of the Chen lab for their discussions and suggestions throughout this study.

## Author contributions

Conceptualization: Qingyang Yin, Liang Chen. Data curation: Qingyang Yin. Formal analysis: Qingyang Yin. Funding acquisition: Liang Chen. Investigation: Qingyang Yin, Liang Chen. Methodology: Qingyang Yin, Liang Chen. Project administration: Liang Chen. Resources: Qingyang Yin, Liang Chen. Software: Qingyang Yin. Supervision: Liang Chen. Validation: Qingyang Yin. Visualization: Qingyang Yin. Writing – original draft: Qingyang Yin. Writing – review & editing: Liang Chen.

## Data availability statement

The codes are available at https://github.com/qyyin0516/xNNDriver-and-xAEDriver. All data from the DepMap database were downloaded from https://depmap.org/portal/download/all/ (version: 23Q2). The COSMIC database can be downloaded at https://cancer.sanger.ac.uk/cosmic/download/cosmic/ (version: v102), and categories of clinical significance for SNVs can be found at https://useast.ensembl.org/info/genome/variation/prediction/predicted_data.html#consequences/. The ClinVar database can be downloaded at https://www.ncbi.nlm.nih.gov/clinvar/, and categories of clinical significance for SNVs can be found at https://www.ncbi.nlm.nih.gov/clinvar/docs/clinsig/. Reactome pathway annotations and hierarchy can be downloaded at https://reactome.org/download-data. The list of 3,008 frequently mutated candidate genes can be found at https://github.com/idekerlab/DrugCell.

