## Supplementary material for "Interpretable neural networks prioritize cancer driver genes from genome-wide dependency landscapes": supplementary information.docx

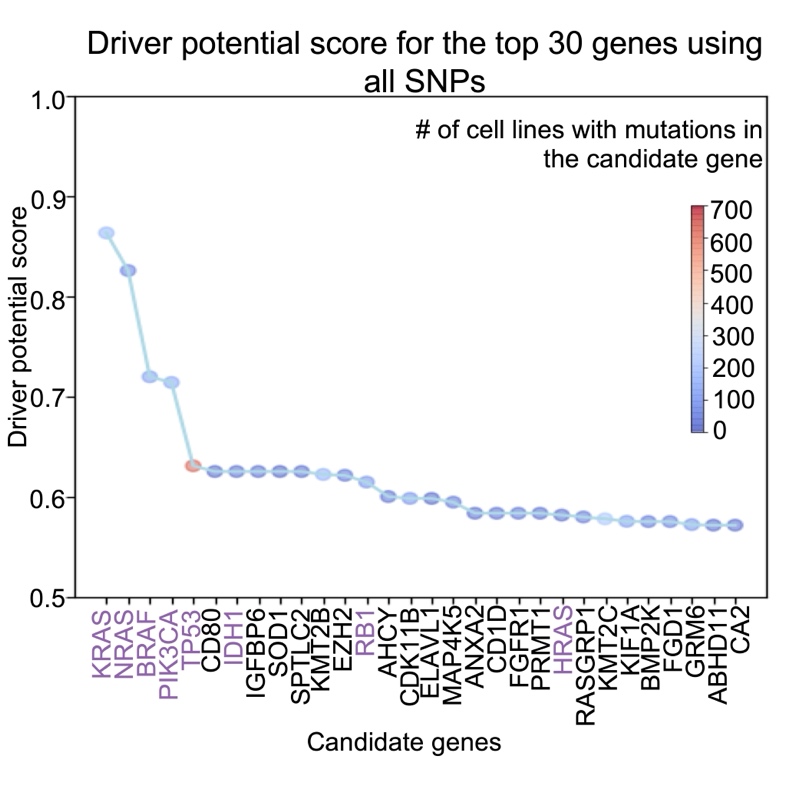

S1 Fig. Prediction performance of xNNDriver for the top 30 genes with the highest driver potential scores using all SNPs. All somatic SNP mutations, no matter whether annotated as functional or not, are used. Data point color indicates the number of cell lines with mutations in that gene. Gene name color distinguishes whether a gene is a known driver or not (purple: driver, black: non-driver), as documented by DepMap.

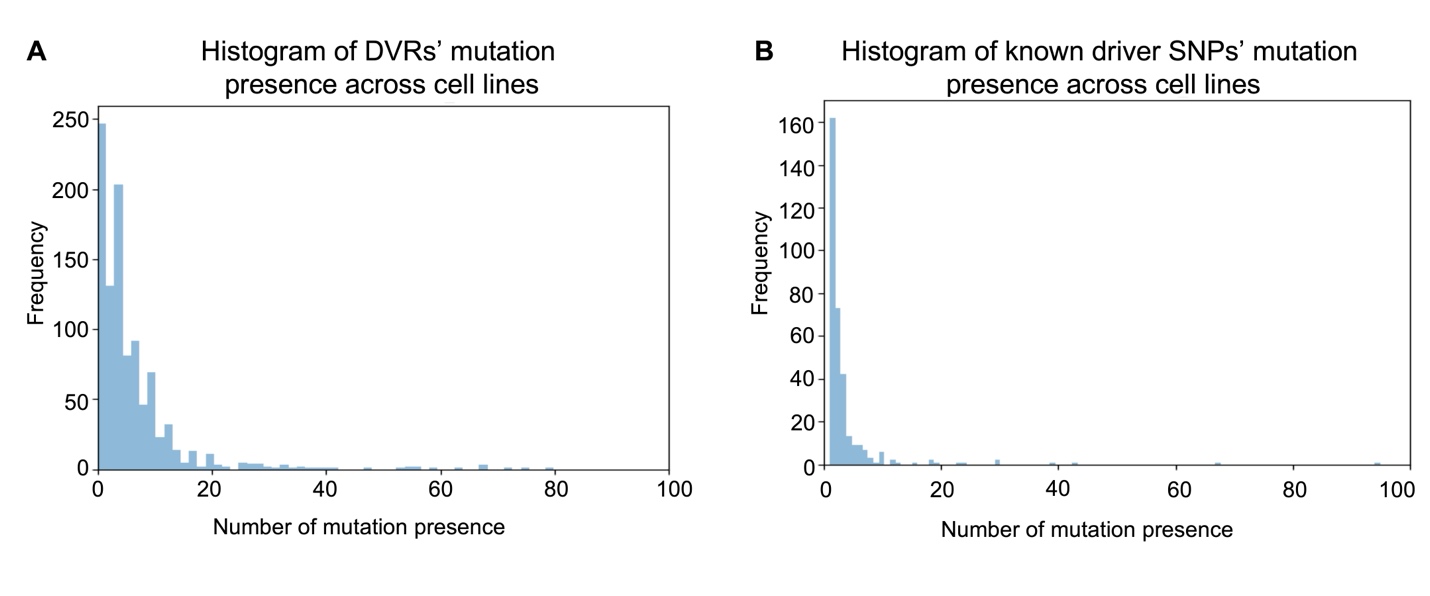
 **S2 Fig. Comparison of mutation frequency distributions between DVRs from xAEDriver and known functional driver SNPs**. The histograms display the frequency of mutation occurrences across cell lines for (**A**) DVRs and (**B**) known driver SNPs.

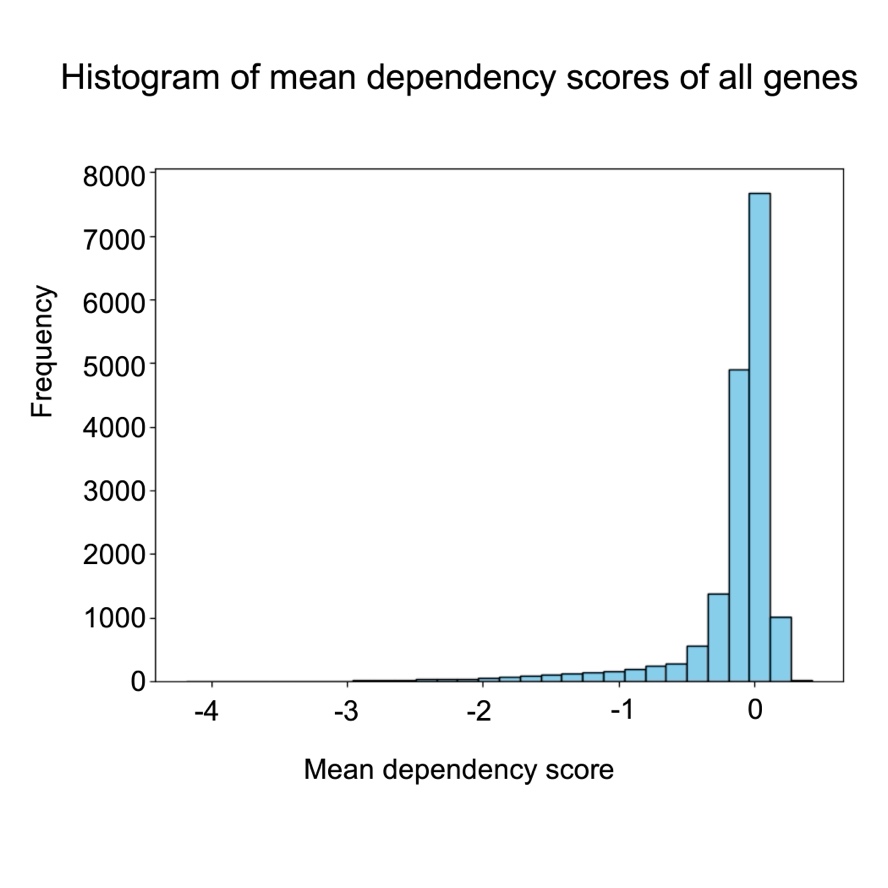

**S3 Fig. Distribution of mean dependency scores of all genes.** The histogram illustrates the distribution of mean dependency scores across all cell lines for all genes in the DepMap gene dependency dataset.

S1 Table. (separate file) Comprehensive results of all xNNDriver models using functional SNPs. Each model corresponds to a frequently mutated candidate gene (3,008 in total). Information includes the following metrics: driver potential score, the number of cell lines with mutation presence, the number of cell lines without mutation presence, and whether the gene is annotated as a driver based on DepMap.

S2 Table. (separate file) Comprehensive results of all xNNDriver models using all SNPs. Each model corresponds to a frequently mutated candidate gene (3,008 in total). Information includes the following metrics: driver potential score, the number of cell lines with mutation presence, the number of cell lines without mutation presence, and whether the gene is a driver based on DepMap.

S3 Table. (separate file) Important pathways identified from xNNDriver for candidate driver genes. This table includes a comprehensive summary combining results for all 15 genes, as well as 15 individual sheets detailing results for each gene. The overall table also provides the adjusted p-values from the Wilcoxon tests on the comparison of coefficients of variation of pathway expression between cell lines with and without mutations.

S4 Table. (separate file) Pathway importance from xAEDriver. This table includes the average $\boldsymbol{R}_{\boldsymbol{p}}$ values across all cell lines. Pathways marked with “_copy1” indicate artificial nodes created to preserve the pathway tree structure.

S5 Table. (separate file) Test set $\boldsymbol{R}^{\boldsymbol{2}}$ values of LASSO regression models based on DVRs and random somatic mutations. Random control results are averaged over 50 runs.

S6 Table. (separate file) Average pathway importance ($\boldsymbol{R}_{\boldsymbol{p}}$ values) and rankings for the DVR-divergent and DVR-convergent groups. Reported metrics include average $\boldsymbol{R}_{\boldsymbol{p}}$ and rank for each group.
